# Six drivers of aging identified among genes differentially expressed with age

**DOI:** 10.1101/2024.08.02.606402

**Authors:** Ariella Coler-Reilly, Zachary Pincus, Erica L. Scheller, Roberto Civitelli

## Abstract

Many studies have compared gene expression in young and old samples to gain insights on aging, the primary risk factor for most major chronic diseases. However, these studies only describe associations, failing to distinguish drivers of aging from compensatory geroprotective responses and incidental downstream effects. Here, we introduce a workflow to characterize the causal effects of differentially expressed genes on lifespan. First, we performed a meta-analysis of 25 gene expression datasets comprising samples of various tissues from healthy, untreated adult mammals (humans, dogs, and rodents) at two distinct ages. We ranked each gene according to the number of distinct datasets in which the gene was differentially expressed with age in a consistent direction. The top age-upregulated genes were TMEM176A, EFEMP1, CP, and HLA-A; the top age-downregulated genes were CA4, SIAH, SPARC, and UQCR10. Second, the effects of the top ranked genes on lifespan were measured by applying post-developmental RNA interference of the corresponding ortholog in the nematode C. elegans (two trials, with roughly 100 animals per genotype per trial). Out of 10 age-upregulated and 9 age-downregulated genes that were tested, two age-upregulated genes (*csp-3*/CASP1 and *spch-2*/RSRC1) and four age-downregulated genes (*C42C1.8*/DIRC2, *ost-1*/SPARC, *fzy-1*/CDC20, and *cah-3*/CA4) produced significant and reproducible lifespan extension. Notably, the data do not suggest that the direction of differential expression with age is predictive of the effect on lifespan. Our study provides novel insight into the relationship between differential gene expression and aging phenotypes, pilots an unbiased workflow that can be easily repeated and expanded, and pinpoints six genes with evolutionarily conserved, causal roles in the aging process for further study.

**Graphical Abstract:** 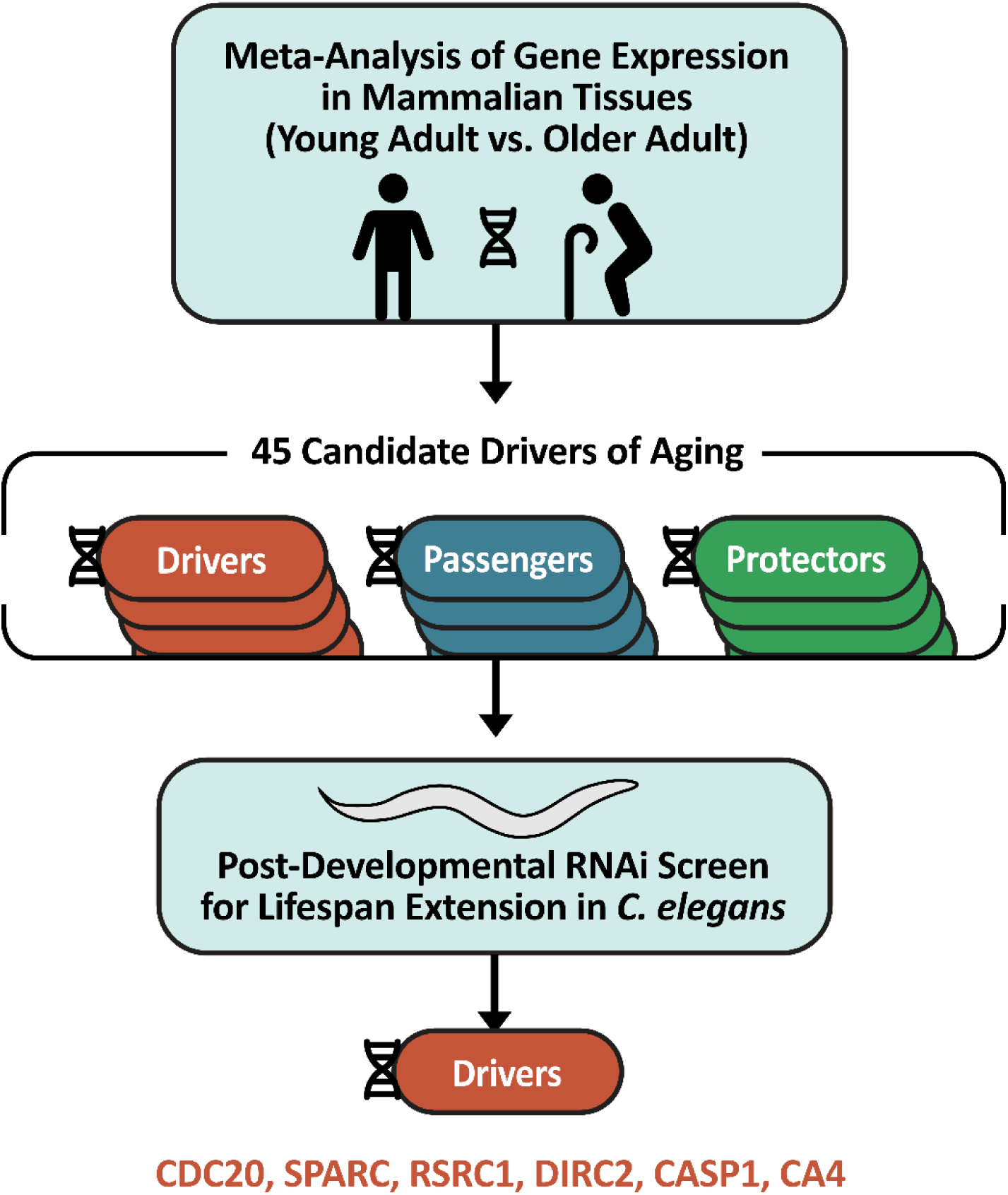

## Introduction

Advanced age is the primary risk factor for most chronic diseases, and as our population ages, the social and economic burden of chronic disease continues to grow year by year [1]. The field of geroscience has emerged to study the mechanisms underlying aging itself and develop strategies to combat age-related decline, or senescence, at the source. According to the widely accepted evolutionary theory of aging, senescence is pervasive because there is negligible selection pressure during the post-reproductive period, a phenomenon known as the “selection shadow” [2, 3]. It is therefore crucial to study the role of genetic variants and gene expression changes in aging in order to uncover potentially advantageous adjustments that have been masked by the selection shadow.

Specialized approaches are needed to detect age-related gene expression signals, which are often subtle and widespread rather than striking and targeted. Indeed, as noted in the *Handbook of the Biology of Aging*, differentially expressed genes (DEGs) with the largest fold changes are frequently found to be downstream targets rather than upstream regulators [4]. Moreover, age-related phenomena such as transcriptional drift create reproducible expression patterns that are nonetheless stochastic and unregulated, further obscuring meaningful signals [5–7]. As stochastic signals are unlikely to replicate across species and tissues, and since frequency is more meaningful than fold change, drivers of aging may ideally be identified using a multi-species, multi-tissue meta-analysis using the value-counting method. This strategy, first pioneered by Magalhães et al. in 2009 [8] and further developed by Palmer et al. in 2021 [9], has been used to catalog numerous individual DEGs as well as broader patterns in functional enrichment and pathway analysis. However, translation of such findings into actionable therapeutic strategies is challenging. Any upregulated gene presumed to be a driver of aging could just as easily be a compensatory geroprotective response or an unimportant downstream effect, often called a “passenger” to contrast with the aforementioned “driver” [5, 10]. In other words, as the age-old adage warns, correlation does not necessarily equal causation.

Functionally evaluating genes related to aging also presents special challenges. Stable cell lines cannot be used to study aging *in vitro* because of the immortal nature of such lines. *In vivo* models are more useful, but require time and resources to age the animals and monitor them until their natural death. In mammals, this can entail years of labor, and this is surely one reason why the short-lived nematode *C. elegans* has been such a popular model organism in geroscience for decades [11]. While nematodes are only distant relatives of humans, they share remarkably similar features of post-reproductive senescence, including sarcopenia and reduced motility, deteriorated learning and memory, and weakened immunity [12–14]. In contrast to humans or any mammal, these age-dependent changes occur on a compressed time-scale of days rather than years, with an average lifespan of only a few weeks. On a genetic level, orthologs of roughly half of all human genes have been identified, and tools have been developed to rapidly, easily, and inexpensively manipulate those genes in *C. elegans* worms, making them an ideal choice for genetic screens [15]. However, it is difficult to substantiate findings in *C. elegans* as relevant to human physiology without any means of contextualizing the results in mammalian systems.

Here, we introduce a workflow to unify two separate approaches, analysis of mammalian DEGs and genetic screening in *C. elegans*, leveraging the strengths of each to mitigate their respective weaknesses. We first performed a meta-analysis comparing gene expression in young adults vs older adults using publicly available datasets comprising samples of various tissues from healthy, untreated mammals (humans, dogs, and rodents). DEGs were ranked by the consistency of differential expression with age across the largest number of datasets. The highest ranking DEGs with known orthologs in *C. elegans* were then tested using post-developmental RNAi lifespan assays. Ultimately, we identified six genes with evolutionarily conserved, causal effects on aging that may be prioritized for future mechanistic studies. In addition, we have established a proof of principle for a unified approach for studying evolutionarily conserved mechanisms of aging.

## Material and Methods

### Meta-Analysis Dataset Selection

This meta-analysis was designed as a simple and scalable approach that is nonetheless highly capable of identifying a collection of genes consistently associated with mammalian aging. The intention was to extricate subtle but meaningful age-related signals from a background of transcriptional drift and stochastic changes that are unlikely to replicate across species, tissues, and experimental platforms. Thus, as the inclusion criteria and exclusion criteria detailed in Table 1 show, any datasets comprising samples from mammals representative of typical individuals at both young adult and older adult time-points were eligible for inclusion.

**Table 1.** Inclusion and exclusion criteria for the meta-analysis of genes differentially expressed during mammalian aging.

| Criteria | Included | Excluded |
| --- | --- | --- |
| Data Availability | Stored in NCBI GEO database,<br>Includes “age” as subset variable | All other datasets |
| Species | Mammal (human, mouse, rat, dog) | Non-Mammal (e.g., drosophila, C. elegans...) |
| Tissue Type | All (heart, muscle, brain, liver, fat,<br>immune cells...) | None |
| Age | Young adult and older adult<br>in distinct groups | Early development, including embryonic stages<br>as well as juveniles |
| Condition | Healthy, untreated | All diseases (e.g. lupus, tumor samples),<br>All interventions including drugs as well as<br>lifestyle interventions like diet and exercise |
| Genotype | Wild-type,<br>No genetic condition specified | Mutant, transgenic animals,<br>Humans with specified genetic conditions |
| Sample Size | Any (no specific minimum) | Datasets with low sample size became naturally<br>excluded when they yielded no DEGs |

**NCBI GEO** = National Center for Biotechnology Information, Gene Expression Omnibus; **DEGs** = Differentially Expressed Genes

Gene expression data were obtained from publicly available datasets hosted on the National Center for Biotechnology Information (NCBI) Gene Expression Omnibus (GEO) repository [16]. The filters “*organism: mammal”* and “*subset variable type: age”* were used to identify roughly 200 curated datasets as candidates for the present study as of March 2021. These datasets were then manually inspected using the inclusion and exclusion criteria (Table 1) to identify 25 suitable datasets, which are listed in Supplementary Table 1.

### Identification of Differentially Expressed Genes (DEGs)

The analysis of gene expression data was conducted using the R software environment version 3.2.3 [17] and a series of packages from the Bioconductor project, including *GEOquery* version 2.40.0 [18], *limma* version 3.26.8 [19], and *BioBase* version 2.30.0 [20]. Briefly, by adapting scripts from GEO’s own GEO2R tool [21], the data was retrieved and translated to R-compatible formats via *GEOquery* and analyzed for DEGs via *limma*. DEGs were calculated by comparing samples from young vs. old tissues for each individual dataset; individual samples were included or excluded from the analysis using the aforementioned inclusion and exclusion criteria (Table 1). To cast a wide net with high sensitivity to detect DEGs even in datasets with small sample sizes, candidate DEGs were identified in individual datasets using the permissive threshold of adjusted p-value < 0.25, controlling for false discovery rates (FDR) using the Benjamini-Hochberg method. Finally, to facilitate analysis across datasets, all DEGs from non-human datasets were converted to their human homologs using the *homologene* package version 1.4.68.19.3.27 [22].

### Value-Counting Method for Ranking DEGs

The genes were then scored using a variation of the value-counting method first established in the cancer field [23] and later applied to age-dependent gene expression [8]. This approach enables the integration of gene expression data from diverse species, tissues, platforms, and experimental designs while remaining highly scalable and reproducible. In brief, genes are ranked by the number of datasets in which they are identified as a DEG according to a chosen threshold. Thus, consistency of differential expression across a variety of datasets is prioritized, while individual effect sizes are discarded.

Here, a new variation of the value-counting method was introduced to further prioritize consistency: ranking was determined based on the absolute value of the difference between upregulation and downregulation scores, where scores were determined by the number of datasets in which the DEG was significantly upregulated and downregulated with age, respectively. This is written formulaically below along with an example.

Let scores *S*_*i*_*Up*__ and *S*_*i*_*Down*__ represent the number of datasets in which gene *i* is significantly upregulated and downregulated with age, respectively. The total score *S*_*i*_ and rank *R*_*i*_ for each gene *i* is calculated as follows:

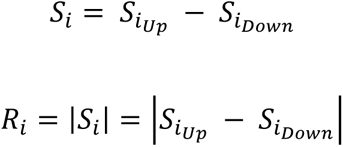

For example, if a gene was significantly (adj. p < 0.25) upregulated in 2 datasets and downregulated in 8 datasets out of the total 25 datasets analyzed, this gene would have a rank of |2 − 8| = 6. It is important to highlight that a gene with a total score and rank of 0 does not necessarily indicate that the gene was differentially expressed in none of the datasets, as it could also be upregulated and downregulated in an equal number of datasets.

Based on the average number of DEGs identified per dataset in the previous step being 1,816 genes out of an average over 30,000 probes per dataset, a binomial distribution with a success rate of 6% and 25 trials can be applied to estimate the final p value for high-ranking genes. For DEGs of rank 6 or above, the cumulative probability *P*(*X* ≥ 6) yields a final p value of 0.003:

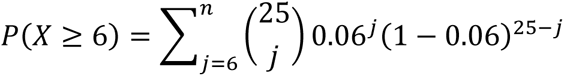

DEGs with a rank of at least 7 (*R*_*i*_ ≥ 7) were further analyzed for gene expression patterns across tissues by normalizing to the number of datasets from each tissue type that were analyzed.

This value-counting analysis was conducted using the python software environment version 3.11.5 [24], and the data were visualized utilizing the *pandas*, *matplotlib*, and *seaborn* packages [25–27].

### Pathway Analysis

DEGs with a rank of at least 6 (*R*_*i*_ ≥ 6) were analyzed using pathway analysis in the R software environment to explore their known roles in key biological processes. The Bioconductor package *clusterProfiler* version 4.8.0 [28] was used to perform gene ontology (GO) enrichment analysis. DEGs were mapped to GO biological processes, cellular components, and molecular functions using standard settings (Benjamini-Hochberg adjusted p < 0.05).

### Identifying Worm Orthologs of DEGs

*C. elegans* orthologs of DEGs with a rank of at least 7 (*R*_*i*_ ≥ 7) were identified using OrthoList 2, which is a compendium of worm genes with human orthologs compiled by a meta-analysis of several orthology prediction methods [29]. Where multiple orthologs were available for a given DEG, the highest confidence ortholog was chosen, as indicated by the number of orthology prediction methods supporting orthology. Where multiple orthologs and/or clones were available for a given gene without any discernible way to prioritize one over another, the first item listed in the results was chosen. The final list of orthologs along with the availability of corresponding RNAi clones is shown in Supplementary Tables 2 and 3. In some cases, when a clone could not be cultured or verified by sequencing (as outlined in the next section below), experiments were conducted using the next clone on the list.

### Worm Culture and Post-Developmental RNAi

Wild-type (N2) *C. elegans* worms were maintained on plates of solid nematode growth media (NGM) seeded with *Escherichia coli* OP50 bacteria at 20°C using standard protocols[30]. *Escherichia coli* HT115 bacteria clones carrying RNAi constructs of interest were obtained from the Ahringer RNAi library [31] and seeded onto solid NGM plates containing Isopropyl β-D-1-thiogalactopyranoside (IPTG) and ampicillin according to standard protocols for the RNAi feeding method [12, 32]. Briefly, for each gene of interest, an individual colony of RNAi bacteria was cultured in liquid LB medium overnight and then seeded onto plates the following day. In parallel, to confirm the identity of the clones, DNA was isolated from the same culture using the QIAprep Spin Miniprep Kit (QIAGEN, Hilden, Germany), and the inserts were sequenced with an M13-forward primer using standard Sanger sequencing services by Azenta Life Sciences (South Plainfield, NJ, USA). The seeded plates were incubated at room temperature for 2-3 days, during which time 2’ fluro-5’ deoxyuridine (FUDR) was added to the plates 24-48 hours before transferring worms. Worms were age-synchronized using the bleaching method with L1 synchronization and allowed to develop to the late L4 stage on standard OP50 plates before being transferred to the plates seeded with the RNAi feeding bacteria, as described in previous post-developmental RNAi screens [33, 34].

### Lifespan Extension Screen

Lifespan assays were conducted using standard protocols [30]. Briefly, worms were scored as alive or dead every two to three days by visual observation: apparently motionless worms were gently prodded with a platinum wire pick, and worms that failed to react were scored as dead and removed from the plate. Worms that left the plate surface and dried on the plate wall were censored, but worms that displayed abnormalities such as internal hatching or vulva rupture were included in all analyses. For the initial screening, the 19 candidate clones were tested across several batches of experiments, with a GFP RNAi negative control group present in every batch, and the well-known *daf-2* RNAi positive control [35] in some selected batches. For every clone tested, the initial screening included roughly 80-100 worms spread across multiple plates (biologic replicates), with approximately 20 worms per plate. For the subsequent validation of the clones that significantly extended lifespan in the initial screening, each group included roughly 100-120 worms, with approximately 25 worms per plate. These were tested in just two batches back-to-back under identical conditions with the same stocks of plates and reagents to minimize batch effects and maximize comparability of results. A GFP RNAi negative control group was included in both batches; the *daf-2* RNAi positive control was included in one batch, while an additional negative control empty L4440 vector RNAi was included in the other batch. To further ensure reproducibility, the N2 worms used during the initial screening and validation experiments were obtained from colonies maintained by separate, independent laboratories (see Acknowledgments for collaborators).

### Lifespan Extension Analysis

Lifespan was defined as the number of days until death, starting from the first day of adulthood (3 days after L1 synchronization). The Online Application for Survival Analysis 2 (OASIS 2) tool was used to calculate mean, median, and maximum lifespans for each group as well as to compare test groups using the log-rank test [36]. An RNAi clone was considered to have extended lifespan if the log-rank test comparing that clone to the GFP RNAi negative control within the same batch was significant (p < 0.05 with Bonferroni multiple test correction) in both the initial screen and the subsequent validation screen. Survival data was then exported and plotted as survival curves using GraphPad Prism version 10.2.3.403 for Windows.

## Results

### Meta-analysis datasets were derived from a variety of mammalian tissues

Twenty-five publicly available gene expression datasets were selected from the NCBI GEO repository according to the inclusion and exclusion criteria outlined in Table 1, and their traits and NCBI identification numbers are listed in Supplementary Table 1. The predominant species represented in this analysis was mouse, constituting roughly half the datasets (13), followed by human (6), then rat (5), then dog (1). Most datasets were derived from muscle (7) and brain (5) tissues, but also well represented were adipose tissues (3) as well as immune cells and their precursors (3), with smaller contributions from the heart, liver, trachea, cochlea, and reproductive tissues (Supp. Fig. 1A). The number of DEGs extracted from each dataset varied widely, ranging from six genes to 3,631 genes (median, 1,509; interquartile range 466 - 3,159). If each instance of a DEG being extracted is considered a data point, then the sum total of data points contributed by most tissues ranged from roughly 4,000 to 11,000; however, cochlea, trachea, and reproductive tissues contributed strikingly fewer, with less than 1,000 data points each (Supp. Fig. 1B). These results reflect the wide variety of studies contributing to this analysis, with varying experimental methods as well as unequal availability of samples from different tissues, particularly from human subjects.

### High-ranking genes were consistently differentially expressed with age across diverse tissues

Using the value-counting method, every gene was assigned upregulation and downregulation scores corresponding to the number of datasets in which that gene was significantly upregulated and downregulated with age, respectively (FDR-adjusted p < 0.25). In general, more genes were commonly upregulated than downregulated with age. Out of a highest possible score of 25 (total number of datasets), the highest downregulation score was 9 (Fig. 1A), and the highest upregulation score was 11 (Fig. 1B). Similarly, only 31 genes achieved a downregulation score of 7 or more, whereas 74 genes reached an upregulation score of 7 or more.

**Figure 1.**
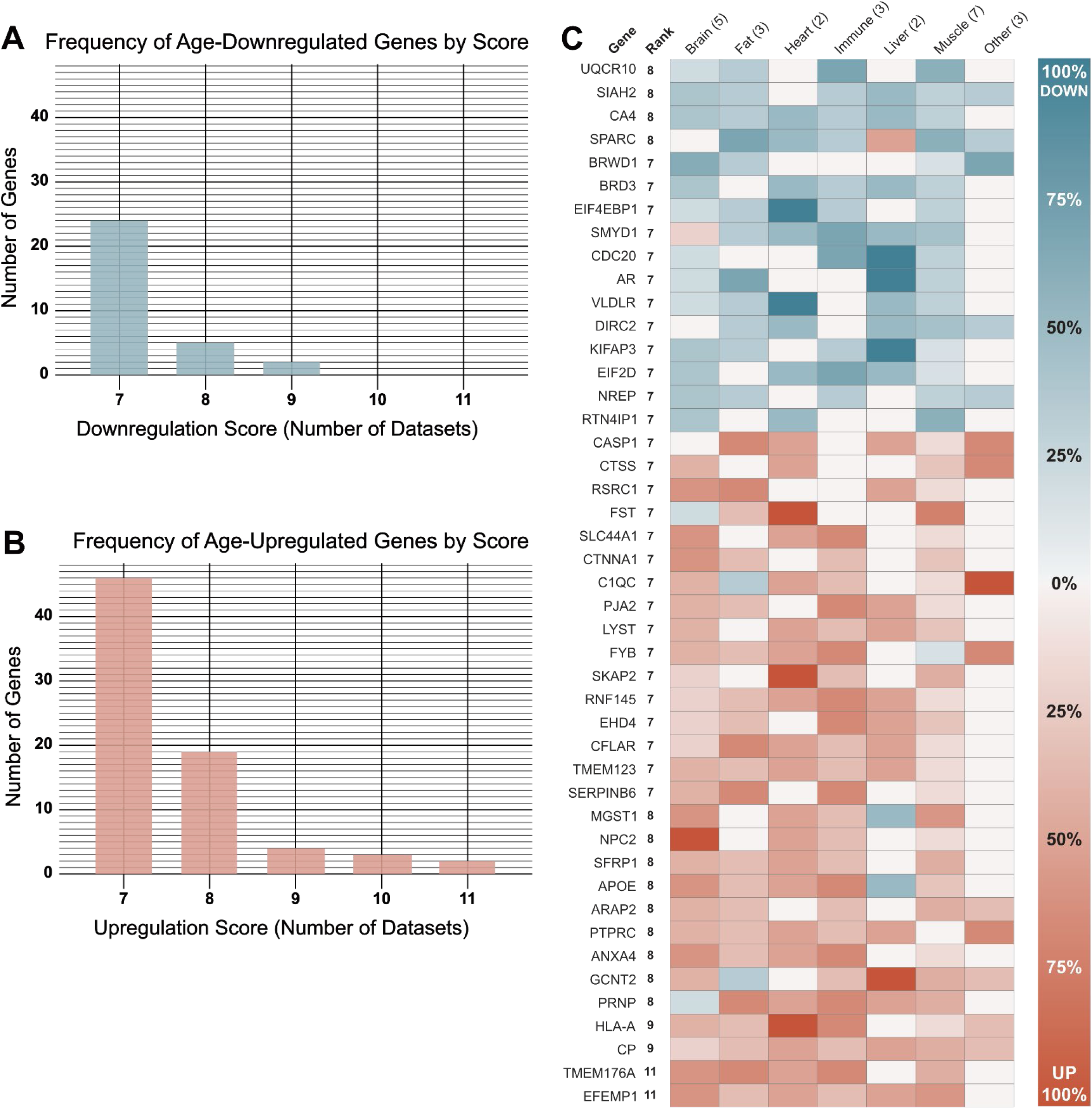
Genes most consistently differentially expressed with age in mammalian tissues. Every gene had a downregulation score *S*_Down_ and an upregulation score *S*_*Up*_ representing the number of datasets in which the gene was significantly downregulated and upregulated with age, respectively (Benjamini-Hochberg adjusted p < 0.25). The rank *R* of each gene was calculated as the absolute value of the difference between these two scores, the total score *S*: *R* = |*S*| = |*S*_*Up*_ − *S*_*Down*_|. (**A**) There were 31 genes with *S*_*Down*_ ≥ 7, and the most consistently downregulated genes had a score of *S*_*Down*_ = 9. (**B**) There were 74 genes with *S*_*Up*_ ≥ 7, and the most consistently upregulated genes had a score of *S*_*Up*_ = 11. (**C**) The heatmap shows the gene symbol, rank *R*, and normalized tissue-specific expression trends for all 45 genes of rank *R* ≥ 7, comprising 29 age-upregulated genes (*S* > 0) and 16 age-downregulated genes (*S* < 0). Heatmap values range from 100% down to 100% up, representing as a percentage the fraction |*S*_*tissue*_/*n*_*tissue*_|, or the total score *S* divided by the number of datasets *n* derived from only the specified tissue type.

To narrow the list of DEGs to those with the most consistent trends, genes were ranked according to the absolute value of the difference between their upregulation and downregulation scores. Thus, genes exhibiting opposing trends in different species or tissues were not ranked highly. While there were 105 genes with a downregulation or upregulation score of at least 7, there were only 45 genes that ranked 7 or above after the opposing score was subtracted. The highest-ranking age-upregulated genes were EFEMP1 (Rank 11), TMEM176A (11), CP (9), and HLA-A (9); the highest-ranking age-downregulated genes were CA4 (8), SIAH2 (8), SPARC (8), and UQCR10 (8). Ranks were used to select DEGs for further analyses and experiments: rank 6 was used as the cut-off to select 130 DEGs for pathway analysis, and rank 7 was used as the cutoff to select 45 DEGs for *in vivo* testing in *C. elegans*.

The 45 highest-ranking DEGs, comprising 16 age-downregulated and 29 age-upregulated genes, are listed in Figure 1C with a heatmap displaying the tissues that contributed to each gene’s rank. For example, EFEMP1, which was tied for the highest-ranking gene, was significantly upregulated in datasets from mouse liver and hematopoietic stem cells, rat heart and adipose tissues, and both human and mouse brain and muscle tissues; EFEMP1 was not significantly downregulated in any of the 25 datasets analyzed. As illustrated in the heatmap, no gene was able to achieve this high rank without being consistently differentially expressed in datasets from at least three distinct tissue types, and often more. The genes CA4 and CP were notable for being consistently differentially expressed across all six major tissue types studied as well as being among the top four highest-ranking downregulated and upregulated genes, respectively.

Excluding tissue types with only one or two datasets, the only gene differentially expressed in 100% of datasets from a given tissue type was NPC2, which was age-upregulated in all five datasets from the brain, as well as handful of datasets from heart, muscle, and immune tissues. Collectively these findings illustrate how the meta-analysis ranking system was able to reveal genes with striking age-related expression patterns.

### Gene ontology patterns were consistent with previous literature

Gene ontology (GO) enrichment analysis was performed to assess how high-ranking DEGs could be categorized into recognizable functional groups and pathways. For this analysis, the cutoff was relaxed to include DEGs of rank 6 and above, yielding a pool of 40 age-downregulated and 90 age-upregulated genes. The GO term matching the largest number of genes from the downregulated pool was the mitochondrial inner membrane, and several additional terms related to mitochondria were enriched as well (Fig. 2A-B). Also strongly represented were both cellular components and molecular functions related to extracellular matrix proteins, particularly collagen. Interestingly, collagen-containing extracellular matrix was also the top GO term for the pool of age-upregulated genes (Fig. 2C-D). However, the overwhelming majority of the GO terms enriched among age-upregulated genes were biological processes related to immune activity, especially adaptive immunity. The results of this pathway analysis largely aligned with expectations and patterns observed in previous studies, reinforcing the validity of the meta-analysis design and execution.

**Figure 2.**
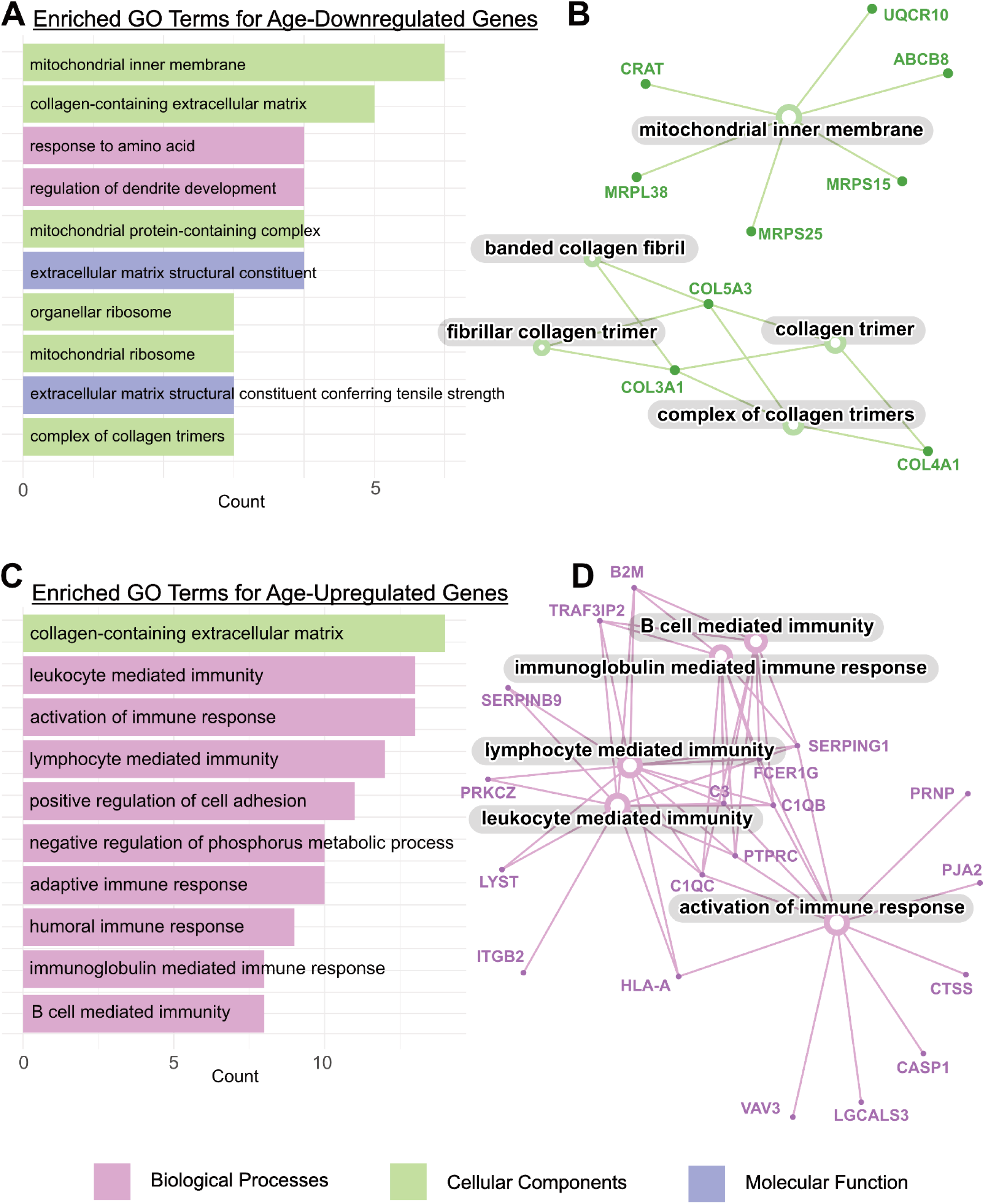
Gene ontology (GO) enrichment analysis of the 130 genes ranked highly for consistent differential expression with age, including 40 age-downregulated genes and 90 age-upregulated genes. For each group, bar charts display the top ten GO terms, colored according to three major GO term categories shown in the key. (**A**) Among age-downregulated genes, the most well-represented GO category was Cellular Components (CC, green). (**B**) The CC category is displayed in more detail in the accompanying network diagram, highlighting the downregulation of collagen-related and mitochondrial membrane-related genes. (**C**) Among age-upregulated genes, the major category was Biological Processes (BP, red). (**D**) The BP category is expanded to show the variety of immune response-related genes upregulated with age. The data were analyzed and visualized in R using the *clusterProfiler* package standard settings including Benjamini-Hochberg adjusted p < 0.05.

### Knocking down orthologs of several mammalian DEGs extended lifespan in *C. elegans*

To prepare for *in vivo* experiments in the model organism *C. elegans*, the database OrthoList 2 was searched for worm orthologs corresponding to the highest-ranking mammalian DEGs. Out of the 45 DEGs queried, 27 (60%) were matched to at least one worm ortholog, including 11 age-downregulated genes (Supp. Table 2) and 16 age-upregulated genes (Supp. Table 3). After excluding orthologs with no corresponding verifiable RNAi clone, a total of 19 genes were available for *in vivo* testing in *C. elegans*, including roughly equal quantities of age-downregulated (9) and age-upregulated (10) genes.

**Table 2.**
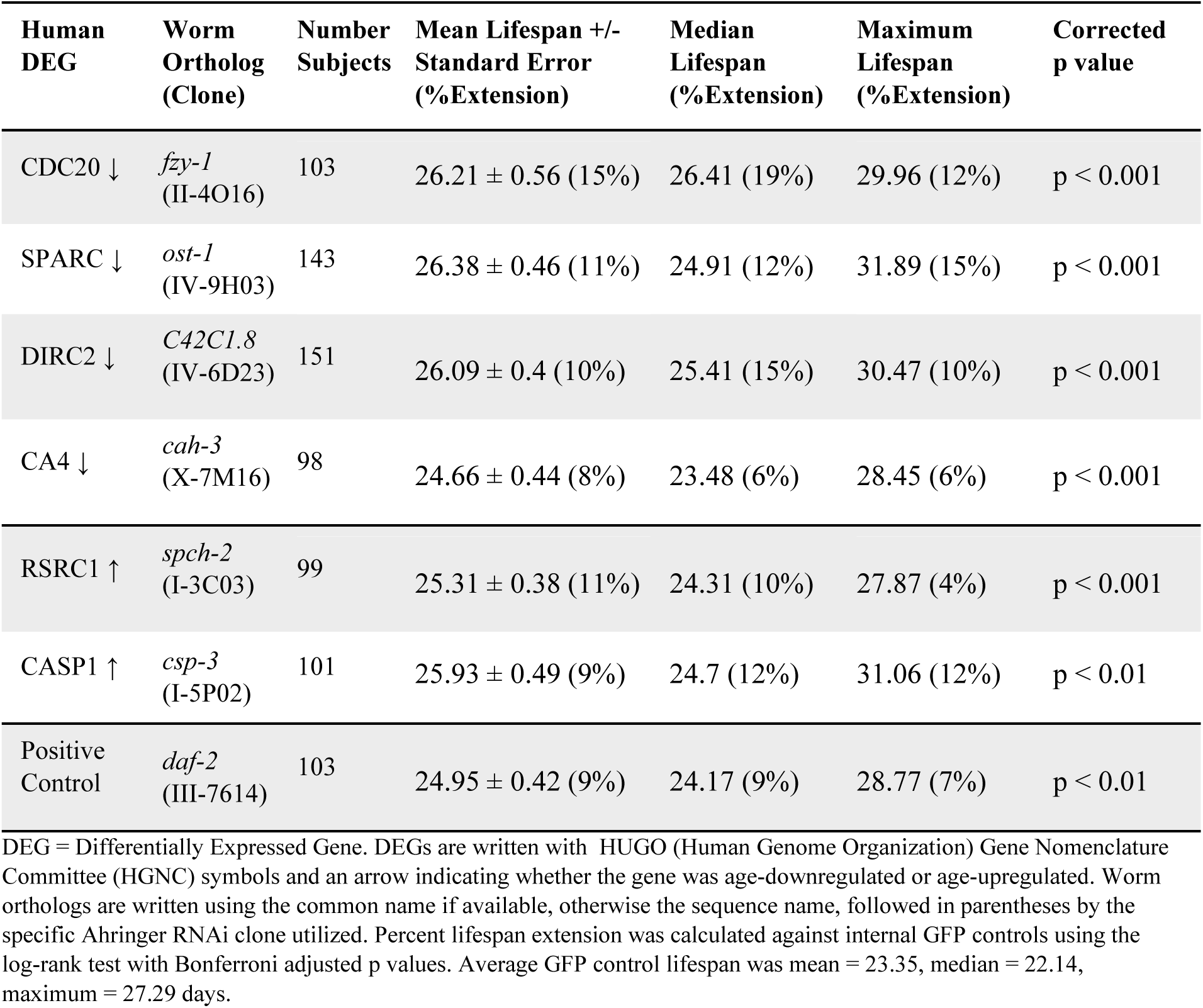
Lifespan extension in *C. elegans* via post-developmental RNAi of orthologs of mammalian DEGs.

To probe the role of each ortholog in senescence while excluding effects on embryonic and juvenile development, bacteria carrying the RNAi clones were fed to the worms post-developmentally. A two-tiered screen was performed, with an initial screening (n ≈ 80 – 100 worms per group) for all 19 orthologs followed by independent validation experiments (n ≈ 100 ‒ 150 worms per group) for RNAi clones that exhibited significant lifespan extension during the screening. Knockdown of six out of 19 (or 32%) of the orthologs of mammalian DEGs significantly extended lifespan in *C. elegans* during both tiers of the screen relative to within-batch GFP controls (log-rank test, Bonferroni-adjusted p < 0.05). In order from largest to smallest mean lifespan extension, these knockdowns targeted: *fzy-1* (ortholog of CDC20), *ost-1* (SPARC), *spch-2* (RSRC1), *C42C1.8* (DIRC2/SLC49A4), *csp-3* (CASP1), and *cah-3* (CA4), which were predominantly (66%) age-downregulated DEGs (Fig. 3A-F). The only two age-upregulated targets were *spch-2* (RSRC1) and *csp-3* (CASP1), while the rest were age-downregulated. Mean lifespan extension ranged from 9% to 15%, median extension from 6% to 19%, and maximum lifespan 4% to 15% relative to within-batch GFP controls, which averaged a lifespan mean of 23.35, median 22.14, and maximum 27.29 days (Table 2). The variability in the validation experiments was very low, as illustrated by the overlapping survival curves for the three independent negative control groups (Fig. 3G). The degree of lifespan extension was comparable to the positive control RNAi against *daf-2* (Fig. 3H), which extended mean lifespan by 9%, median by 9%, and maximum by 7% (n = 103, log-rank test, Bonferroni-adjusted p < 0.01, Table 2).

**Figure 3.**
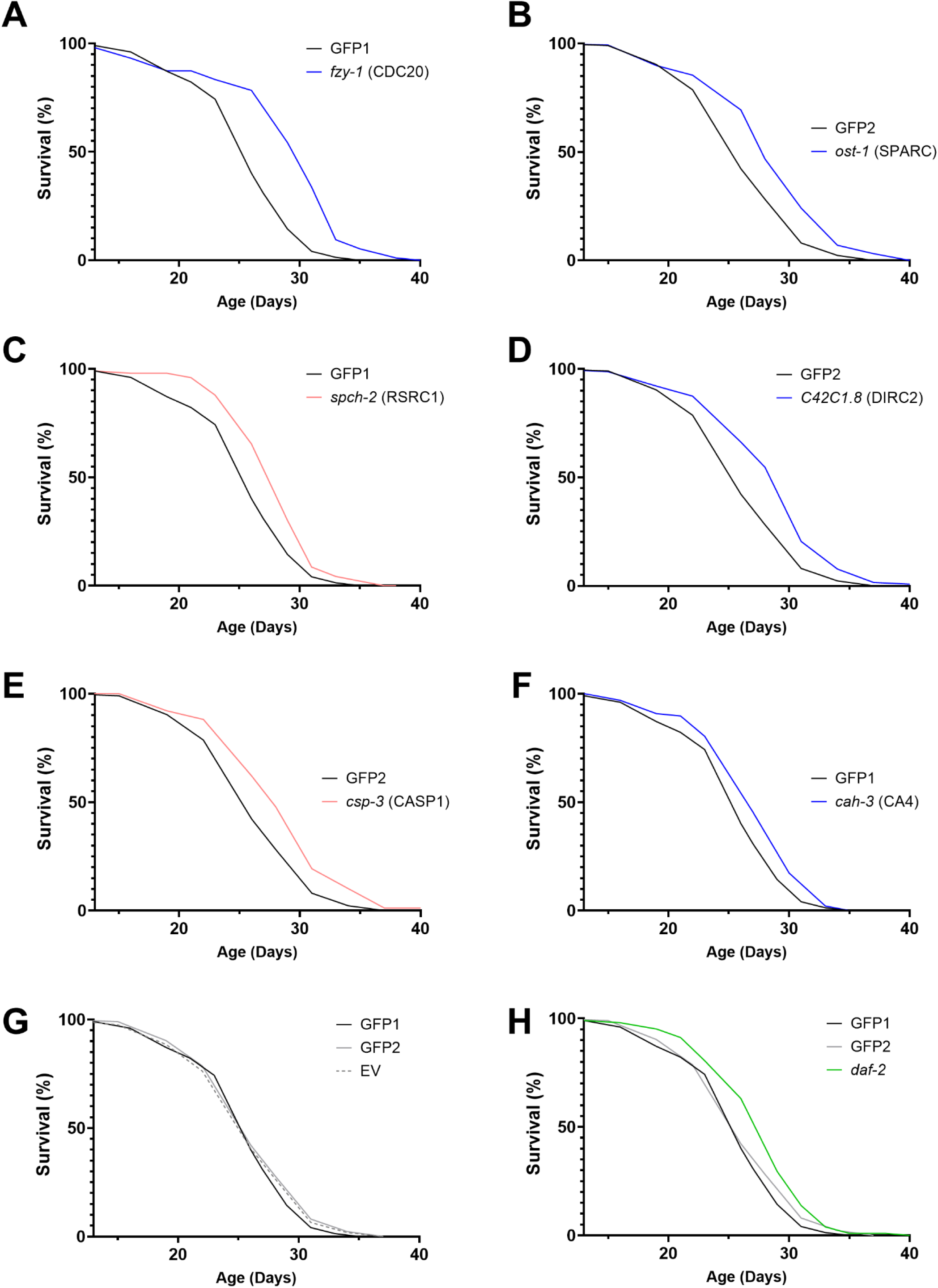
*C. elegans* survival curves for lifespan-extending post-developmental RNAi targeting orthologs of mammalian DEGs, including age-downregulated (blue) and age-upregulated (red) genes. Each RNAi clone significantly extended lifespan (**A**-**F**, log-rank test, Bonferroni-adjusted p < 0.05) in two independent experiments, and the results of the second experiment are shown here (n ≈ 100 ‒ 150 worms per group). This experiment was performed in two back-to-back batches with an internal GFP control group in each batch (GFP1 for batch 1, GFP2 for batch 2). (**G**) Negative control GFP groups performed very similarly to each other (solid lines) and to the alternative empty L4440 vector control (dotted lines). (**H**) The RNAi clone Ahringer III-7614 against *daf-2* was used a positive control (green). For detailed quantification, see Table 2.

Lastly, the expression patterns of the six genes that extended lifespan in worms were re-examined in our mammalian datasets (Supp. Fig. 2). CASP1 and RSCR1 were age-upregulated in seven datasets each with no age-downregulation. CA4, DIRC2, and CDC20 were age-downregulated in seven or eight datasets each with no age-upregulation. SPARC was age-downregulated in nine datasets but also age-upregulated in one dataset derived from mouse liver. All six were differentially expressed with age in multiple mouse tissues; all except CDC20 in human tissues; and all except CA4 in rat tissues. All six genes were differentially expressed in at least one of the seven datasets from muscle, which is unsurprising, but also one of only two datasets from liver, which is a much higher rate. Expression patterns in adipose tissue were also prominent: CASP1, RSRC1, and SPARC were differentially expressed in two of three fat datasets, and CA4 and DIRC2 in one dataset each. Finally, RSRC1, CA4, and CDC20 were differentially expressed in certain brain tissues including human frontal cortex and mouse neocortex and striatum. In summary, the most striking pattern was that knockdown of age-upegulated DEGs were not more likely than age-downregulated DEGs to extend lifespan, and the prominent contributions from adipose and liver tissue were also notable.

## Discussion

This study has established a geroscience-specific workflow to channel large quantities of gene expression data into a streamlined list of actionable targets using accessible, scalable tools in computational biology and *C. elegans* research. The goal of this approach is to maximize the value of existing research by harnessing these readily available datasets and methods effectively to produce novel, valuable discoveries.

Over the past few decades there have been copious studies comparing gene expression in tissues from older versus younger subjects in a variety of species [37, 38], and these generally culminate in conclusions based on functional enrichment analysis. In general, advanced age has been associated with upregulation of immune and inflammatory pathways but downregulation of the electron transport chain and other mitochondrial activities as well as collagen and other extracellular matrix proteins [8, 37, 39], and our results were consistent with these established trends. However, therapeutic directions cannot be extrapolated from purely observational gene expression data, where drivers of aging cannot be distinguished from compensatory protective responses and irrelevant downstream effects. Moreover, there is no guarantee that functional groups reflect concerted biological activities, and they are biased in favor of well-defined gene-sets, which are both important limitations to consider.

Two of our highest-ranking individual genes, EFEMP1 (Rank 11) and CP (Rank 9), which were consistently age-upregulated over all six major tissue types, have known associations with age-related pathologies, and have also been classified as age-associated in previous similar meta-analyses [8, 9]. EFEMP1, also known as fibulin-3, is an extracellular matrix glycoprotein strongly associated with aging pathologies: overexpression contributes to age-related macular degeneration, high plasma levels are associated with signs of brain aging and higher risk of dementia, and upregulation of this gene is associated with Werner syndrome, a premature aging condition [40, 41]. On the other hand, CP, or ceruloplasmin, is a copper-binding glycoprotein involved in iron metabolism and defense against oxidative stress; decreased CP activity is associated with advanced age and age-related diseases, such as Parkinson’s and Alzheimer’s disease [42–44]. From this context, we may infer that although both genes exhibit similar expression profiles, EFEMP1 likely plays a role driving age-related pathology, whereas CP may be upregulated with age as a compensatory response to amplify its protective effects. However, even for well documented genes like these, such inferences still involve speculation, and there are many other DEGs that are much less clearly characterized without supplemental information.

We focused on funneling our DEGs into a *C. elegans* RNAi lifespan screen to gain insights on the role of each gene in aging and longevity. Two of the ten tested age-<u>up</u>regulated genes extended lifespan when knocked down in *C. elegans*: *csp-3*,an ortholog of CASP1, and *spch-2*, an ortholog of RSRC1. Caspases are proteases involved in apoptosis and inflammation [45], and CASP-1 is particularly well known a major component of the NLRP3-CASP1 inflammasome and a promising therapeutic target for Hutchinson-Gilford progeria, another premature aging syndrome, and Alzheimer’s disease [46, 47]. In fact, pharmacological CASP-1 inhibitors have demonstrated to be protective against cognitive decline in mouse models of Alzheimer dementia [48, 49]. Interestingly, CASP1 was not differentially expressed in any of the brain datasets we examined, which included samples from human frontal cortex. However, there is evidence that CASP1 is overexpressed in the frontal cortex and hippocampus of patients with Alzheimer’s disease [47].The novel discovery that RNAi inhibition of an orthologous caspase extends lifespan in worms may suggest an evolutionarily conserved role for caspases in driving inflammaging and age-related neurodegeneration.

RSRC1, on the other hand, is much less well studied, but appears to have a role in brain development: RSRC1 polymorphism is associated with schizophrenia [50], and patients homozygous for loss-of-function RSRC1 mutations exhibit developmental delay and intellectual disability [51, 52]. In our meta-analysis, RSRC1 was age-upregulated in three of the five datasets from brain tissue, and post-developmental knockdown produced lifespan extension. Collectively, these results suggest that RSRC1 is crucial for early development but functions aberrantly late in life as a driver of aging.

Four of the nine tested age-<u>down</u>regulated genes we tested extended lifespan when knocked down in *C. elegans*, including orthologs of two of the four highest-ranking (Rank 8) age-downregulated DEGs: *ost-1*, ortholog of SPARC; and *cah-3*, an ortholog of CA4. SPARC, also known as osteonectin, is a highly conserved extracellular matrix glycoprotein; it is expressed ubiquitously, though primarily in adipocytes [53, 54]. It also plays a role in bone development and turnover and wound healing, especially in corneal tissue [55–57]. In adipocytes, SPARC has been linked to adipose fibrosis, age-related inflammation and metabolic dysfunction, as well as diabetes and its complications (nephropathy and retinopathy), and obesity [54, 58, 59]. In our meta-analysis, SPARC was downregulated with age in all major tissues studied except the brain and liver; the most pronounced pattern was in adipose tissue, where expression decreased with age in two of the three datasets. Interestingly, SPARC is also associated with liver fibrosis [60], and it was in fact upregulated in our dataset from mouse liver. The large amount of available literature underscores SPARC’s pleiotropic functions and open questions about its role in aging. Our work revealed tissue-specific changes in SPARC expression with age and the discovery that post-developmental SPARC knockdown extends lifespan in *C. elegans*.

Carbonic anhydrases are crucial for such fundamental biological processes as regulating pH and transporting carbon dioxide, and they are ubiquitous from microbes to mammals [61]. We found one member of this enzyme family, CA4, to be consistently downregulated with age across all six major tissue types studied, including human and mouse brain datasets. In humans, CA4 mutations contribute to retinitis pigmentosa [62], but in rodents, CA4 has also been studied as a mechanism for extracellular buffering in the brain [63, 64]. Carbonic anhydrase inhibitors have been pursued as potential treatments for brain edema, glaucoma, epilepsy, cancer, and mountain sickness [61]. Here we showed, for the first time, that downregulating the CA4 ortholog, *cah-3*, post-developmentally extended lifespan in *C. elegans*, suggesting that less CA4 activity may be needed in late life.

The largest lifespan extension achieved in our study was via RNAi knockdown of the CDC20 ortholog *fzy-1*, a result consistent with a previous study (Xue et a. 2007). Cell division cycle 20 (CDC20) is an evolutionarily conserved, positive regulator of cell division essential for life in both worms and mammals [65, 66]. The concentration and activity of CDC20 must be tightly regulated, as hyperactivity is associated with aneuploidy and oncogenesis [67–69], and downregulation is associated with premature cellular senescence [70]. Interestingly, although adult *C. elegans* are post-mitotic creatures, genes associated with cell proliferation and differentiation modulate worm lifespan through mechanisms that are thought to be evolutionarily conserved [71]. We found that CDC20 was most downregulated in mouse dendritic cells, hematopoietic stem cells, and rodent liver tissues, which may reflect a decline in proliferation of those tissues.

Lastly, we found that post-developmental knockdown of the DIRC2 ortholog *C42C1.8* extends lifespan in *C. elegans*. DIRC2, or disrupted in renal carcinoma 2, is also known as solute carrier family 49 member 4, or SLC49A4. Aside from the eponymous roles in renal carcinoma and solute transport (specifically, lysosomal export of vitamin B6), very little is known about this gene [72], or the worm ortholog *C42C1.8,* which has no common name. Our work demonstrates that DIRC2 is actually one of the most consistently age-downregulated genes in mammals, with expression declining in both human and mouse muscle tissues and rodent fat, heart, liver, and trachea. The example of DIRC2 demonstrates the power of our approach to identify promising, understudied targets for further investigation.

*C. elegans* has long been used to investigate the mechanisms of aging using well developed functional genomics tools. There were two genome-wide RNAi longevity screens, by the Ruvkun [73] and Kenyon [74] groups, each boasting 70-80% coverage of all open-reading frames. Due to very high false negative rates, they identified a combined total of 120 longevity genes, with only four genes in common [75]. The Ruvkun group performed a follow-up screen using post-developmental instead of embryonic RNAi on 2,700 genes essential for development [33], and similar smaller studies have been published by others since [34, 76]. The yield of lifespan extending gene activations out of total genes tested was less than 1% for genome-wide screens, and 2.4% for the post-developmental screen of essential genes, reflecting the importance of antagonistic pleiotropy in aging [3, 77]. A more recent study achieved a yield of 44% when testing orthologs of genes differentially expressed with age in human blood, and this study also reported a background rate of 7% yield for randomly chosen genes [78]. Yield is highly dependent on experimental methods such as number of animals and time-points as well as environmental factors like temperature; in this latter case, the authors also tested several of genes under two different conditions (pre- and post-developmentally), raising the yield. Here we reported a yield of 32%, suggesting that roughly one-third of the candidates identified in our meta-analysis are drivers of aging; the remaining two-thirds may have negligible or protective effects, or they may also be drivers of aging but under conditions not tested in this experiment.

There are important limitations to our study, many of which pertain to the nature of RNAi screens and the challenges of modeling human physiology in worms. First, only gene inactivation, not overexpression, could be tested, so we could identify drivers of aging but not geroprotective genes. Secondly, only some of the high-ranking DEGs corresponded to verifiable worm orthologs and were able to be tested, and even those orthologs were selected with varying levels of confidence and specificity. Thirdly, the evolutionary distance between humans and worms limits our interpretation of the functional roles of each gene product, as molecules with very similar structures can play very different biological roles in such distinct species.

It should also be noted that the datasets included in our meta-analysis were derived from neither a complete nor an even distribution of tissue types; for example, there were no datasets derived from the kidneys or intestines, whereas muscle was highly represented. As more datasets are made available, we expect this approach to provide increasing contributions to the geroscience literature. Consistent with this goal, the methods described herein are intended to be accessible and flexible enough for others to reproduce and expand our workflow in future studies. Only basic coding skills in R and python are required to reproduce the meta-analysis, and the GEO2R toolset at the core of our scripts has recently been updated to accept both microarray and RNAseq datasets.

In conclusion, the overall trends we observed in our meta-analysis were consistent with previous literature, but our novel workflow identified six genes with evolutionarily conserved, causal roles in the aging process. Of note, knocking down age-upregulated genes was not more likely to produce life extension than interfering with age-downregulated genes. Thus, our results do not support the commonly held assumption that reversing any changes in age-related gene expression is beneficial, and future studies should further investigate this trend.

## Author Contributions

Conceptualization and Methodology: A. Coler-Reilly, Z. Pincus; Formal analysis and investigation: A. Coler-Reilly, E. Scheller, R. Civitelli; Writing - original draft preparation: A. Coler-Reilly; Writing - review and editing: Z. Pincus, E. Scheller, R. Civitelli; Funding acquisition: A. Coler-Reilly, R. Civitelli; Resources: Z. Pincus, R. Civitelli; Supervision: E. Scheller, R. Civitelli.

## Acknowledgments

This work was supported by grants T32 AR060719, National Institute of Arthritis and Musculoskeletal and Skin Diseases (Skeletal Disorders Training Program), T32 GM007200, National Institute of General Medical Sciences, National Research Science Award (Medical Scientist Training Program), and funds from the Barnes-Jewish Hospital Foundation (B5064-80). We also thank Dr. Eric Greer for providing wild-type (N2) *C. elegans* worms and Dr. Tim Schedl for providing plates of solid nematode growth media (NGM) seeded with *Escherichia coli* OP50 bacteria as well as NGM plates containing Isopropyl β-D-1-thiogalactopyranoside (IPTG) and ampicillin. We also thank Dr. Timothy Peterson for helping with the design of the meta-analysis methods.

## Data Availability

All raw scripts are permanently stored and available for review at the following repository: https://github.com/AriellaStudies/Aging-DEGs

