## Supplemental Materials for "Six drivers of aging identified among genes differentially expressed with age"

**Supplementary Table 1.** Datasets included in the meta-analysis of genes differentially expressed during mammalian aging. GEO ID is an acryonym for Gene Expression Omnibus Identifier, a unique identifier for every data series. Platform refers to the unique identifier for the array used to produce the indicated data series. Samples were broadly categorized into general tissue types for grouped analyses, and any further details were noted in the Tissue Subtype column, where NA = Not Applicable.

| GEO ID | Organism | Platform | Tissue Type | Tissue Subtype |
|---|---|---|---|---|
| GSE71868 | Mus musculus | GPL6885 | Immune | Pulmonary CDC11c+ Cells |
| GSE50821 | Mus musculus | GPL1261 | Muscle | Purified skeletal muscle satellite cells |
| GSE53890 | Homo sapiens | GPL570 | Brain | Frontal cortex |
| GSE55162 | Mus musculus | GPL1261 | Trachea | NA |
| GSE46646 | Mus musculus | GPL1261 | Liver | NA |
| GSE49543 | Mus musculus | GPL339 | Cochlea | NA |
| GSE38718 | Homo sapiens | GPL570 | Muscle | Skeletal muscle |
| GSE28422 | Homo sapiens | GPL570 | Muscle | Skeletal muscle (vastus lateralis) |
| GSE28392 | Homo sapiens | GPL570 | Muscle | Vastus lateralis: type 1 (slow) fibers |
| GSE25941 | Homo sapiens | GPL570 | Muscle | Skeletal Muscle (Vastus Lateralis) |
| GSE25905 | Mus musculus | GPL6246 | Fat | Bone marrow adipocytes |
| GSE25905 | Mus musculus | GPL6246 | Fat | Peripheral adipocytes |
| GSE32719 | Homo sapiens | GPL570 | Immune | BM-HSCs[2] |
| GSE27686 | Mus musculus | GPL1261 | Immune | HSCs[2] |
| GSE24515 | Rattus norvegicus | GPL1355 | Brain | Parietal cortex |
| GSE19677 | Mus musculus | GPL1261 | Brain | Striatum |
| GSE9990 | Rattus norvegicus | GPL341 | Brain | Hippocampus |
| GSE12502 | Canis lupus | GPL3979 | Muscle | Skeletal muscle: biceps femoris |
| GSE11667 | Mus musculus | GPL1261 | Reproduction | Oocytes |
| GSE6718[1] | Rattus norvegicus | GPL1355 | Heart | NA |
| GSE6718[1] | Rattus norvegicus | GPL1355 | Fat | White adipose tissue |
| GSE8150 | Mus musculus | GPL1261 | Brain | Neocortex |
| GSE8146 | Mus musculus | GPL81 | Heart | NA |
| GSE4270 | Rattus norvegicus | GPL890 | Liver | NA |
| GSE6323 | Mus musculus | GPL339 | Muscle | Skeletal muscle: gastrocnemius |

[1]As the data series GSE6718 included two sets of samples, each from a distinct tissue type (heart tissue and white adipose tissue), this series was analzed as two separate datasets and therefore listed in two separate rows. [2]HSCs = hematopoietic stem cells; BM-HSCs = HSCs derived from bone marrow.

**Supplementary Table 2.** Experimental outcomes of the 16 highest-ranking age-downregulated mammalian DEGs[1] in *C. elegans*. Higher rank[2] indicates a DEG was more consistently downregulated with age. The selected *C. elegans* orthologs are designated using the sequence name (Worm Gene), and also noted are the common name (Name), the number of OrthoList2 programs identifying the ortholog noted (Match), and the location of corresponding RNAi clones in the Ahringer library. Genes were excluded from lifespan experiments if there was no known ortholog or if the available RNAi clone(s) failed to grow in standard culture conditions or presented unexpected results upon Sanger sequencing (Exclusion); otherwise, the results of the two-tiered lifespan assays are described as either significantly extending lifespan (Lifespan ↑) or not ( – ).

| Human DEG[1] | Rank[2] | Worm Gene | Name | Match | Ahringer RNAi | Exclusion | Screening | Validation |
|---|---|---|---|---|---|---|---|---|
| **CA4** | 8 | *K05G3.3* | *cah-3* | 1 | X-7M16 | | Lifespan ↑ | **Lifespan ↑** |
| **SIAH2** | 8 | *Y37E11AR.2* | *siah-1* | 4 | IV-8K18 | Failed culture | | |
| **SPARC** | 8 | *C44B12.2* | *ost-1* | 5 | IV-9H03 | | Lifespan ↑ | **Lifespan ↑** |
| **UQCR10** | 8 | | | | | No ortholog | | |
| **AR** | 7 | | | | | No ortholog | | |
| **BRD3** | 7 | *Y119C1B.8* | *bet-1* | 5 | I-9E23 | | Lifespan ↑ | – |
| **BRWD1** | 7 | | | | | No ortholog | | |
| **CDC20** | 7 | *ZK177.6* | *fzy-1* | 1 | II-4O16\|II-4O18 | | Lifespan ↑ | **Lifespan ↑** |
| **DIRC2** | 7 | *C42C1.8* | | 1 | IV-6D23 | | Lifespan ↑ | **Lifespan ↑** |
| **EIF2D** | 7 | *C25H3.4* | | 6 | II-10E20\|II-4H08 | | – | |
| **EIF4EBP1** | 7 | | | | | No ortholog | | |
| **KIFAP3** | 7 | *F08F8.3* | *kap-1* | 6 | III-3N14 | | – | |
| **NREP** | 7 | | | | | No ortholog | | |
| **RTN4IP1** | 7 | *F56H1.6* | *rad-8* | 6 | I-9F05 | | – | |
| **SMYD1** | 7 | *T22A3.4* | *set-18* | 3 | I-5G16 | | – | |
| **VLDLR** | 7 | *T13C2.6* | | 6 | II-5A06 | Failed culture | | |

[1]DEG = Differentially expressed gene, here written using the HUGO (Human Genome Organization) Gene Nomenclature Committee (HGNC) symbol. [2]Rank $R$ was calculated as the absolute value of the difference between upregulation and downregulation scores using the following formula as detailed in the Methods section: $R = |S| = |S_{Up} - S_{Down}|$.

**Supplementary Table 3.** Experimental outcomes of the 29 highest-ranking age-upregulated mammalian DEGs[1] in *C. elegans*. Higher rank[2] indicates a DEG was more consistently upregulated with age. The selected *C. elegans* orthologs are designated using the sequence name (Worm Gene), and also noted are the common name (Name), the number of OrthoList2 programs identifying the ortholog noted (Match), and the location of corresponding RNAi clones in the Ahringer library. Genes were excluded from lifespan experiments if there was no known ortholog or if the available RNAi clone(s) failed to grow in standard culture conditions or presented unexpected results upon Sanger sequencing (Exclusion); otherwise, the results of the two-tiered lifespan assays are described as either significantly extending lifespan (Lifespan ↑) or not ( – ).

| Human DEG | Rank | Worm Gene | Name | Match | Ahringer RNAi | Exclusion | Screening | Validation |
|---|---|---|---|---|---|---|---|---|
| **EFEMP1** | 11 | *F56H11.1* | *fbl-1* | 1 | IV-4N18\|IV-4P14 | | Lifespan ↑ | – |
| **TMEM176A** | 11 | | | | | No ortholog | | |
| **CP** | 9 | | | | | No ortholog | | |
| **HLA-A** | 9 | | | | | No ortholog | | |
| **ANXA4** | 8 | *T07C4.9* | *nex-2* | 3 | III-5F15\|III-8O24 | | – | |
| **APOE** | 8 | | | | | No ortholog | | |
| **ARAP2** | 8 | *F23H11.4* | | 3 | III-1G04 | | – | |
| **GCNT2** | 8 | *T15D6.2* | *gly-16* | 4 | I-6A08 | | – | |
| **MGST1** | 8 | | | | | No ortholog | | |
| **NPC2** | 8 | *R148.6* | *heh-1* | 1 | III-1P05 | Failed Sanger | | |
| **PRNP** | 8 | | | | | No ortholog | | |
| **PTPRC** | 8 | *F56D1.4* | *clr-1* | 1 | II-4M08 | Failed Sanger | | |
| **SFRP1** | 8 | *Y73B6BL.21* | *sfrp-1* | 4 | | No clone | | |
| **C1QC** | 7 | | | | | No ortholog | | |
| **CASP1** | 7 | *Y47H9C.6* | *csp-3* | 1 | I-5P02 | | Lifespan ↑ | **Lifespan ↑** |
| **CFLAR** | 7 | | | | | No ortholog | | |
| **CTNNA1** | 7 | *R13H4.4* | *hmp-1* | 5 | | No clone | | |
| **CTSS** | 7 | *T03E6.7* | *cpl-1* | 2 | V-11I07 | Failed culture | | |
| **EHD4** | 7 | *W06H8.1* | *rme-1* | 4 | V-14F01 | | Lifespan ↑ | – |
| **FST** | 7 | | | | | No ortholog | | |
| **FYB** | 7 | | | | | No ortholog | | |
| **LYST** | 7 | *T01H10.8* | *lyst-1* | 1 | X-5F04 | Failed culture | | |
| **PJA2** | 7 | *Y54E10BR.3* | | 1 | I-7N18 | | – | |
| **RNF145** | 7 | *Y119C1B.5* | | 5 | I-9E22 | | – | |
| **RSRC1** | 7 | *C10G11.9* | *spch-2* | 1 | I-3C03 | | Lifespan ↑ | **Lifespan ↑** |
| **SERPINB6** | 7 | *F20D6.4* | *srp-7* | 3 | V-14F18 | | – | |
| **SKAP2** | 7 | | | | | No ortholog | | |
| **SLC44A1** | 7 | | | | | No ortholog | | |

[1]DEG = Differentially expressed gene, here written using the HUGO (Human Genome Organization) Gene Nomenclature Committee (HGNC) symbol. [2]Rank $R$ was calculated as the absolute value of the difference between upregulation and downregulation scores using the following formula as detailed in the Methods section: $R = |S| = |S_{Up} - S_{Down}|$.

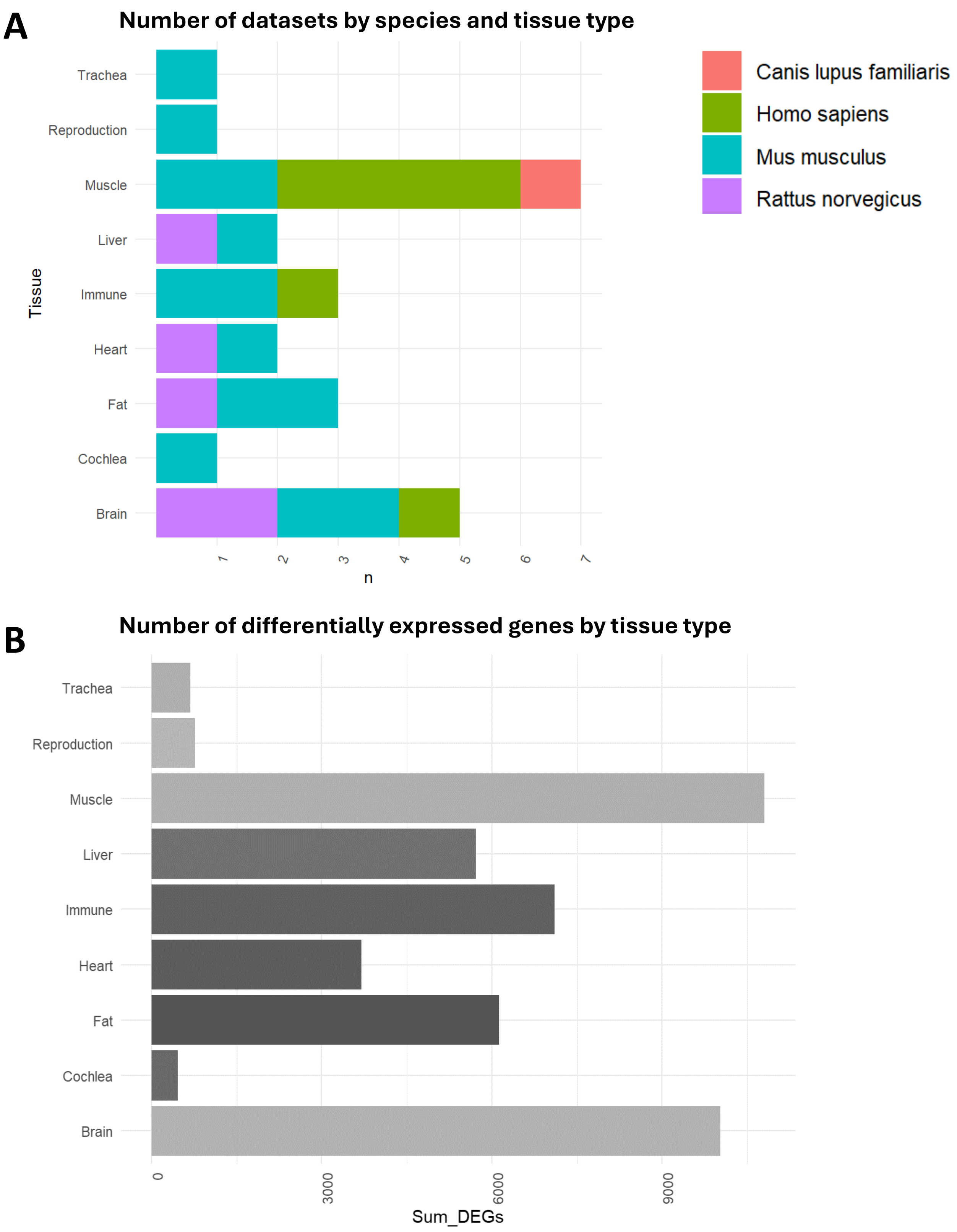


**Supplementary Figure 1.** The 25 datasets studied were derived from an uneven distribution of species and tissues, and the resulting DEGs were also unevenly distributed across tissue types. (**A**) The total number of datasets (n) derived from each major tissue type, color-coded by the species of origin. (**B**) The sum total number of instances DEGs were identified across all datasets derived from each major tissue type.

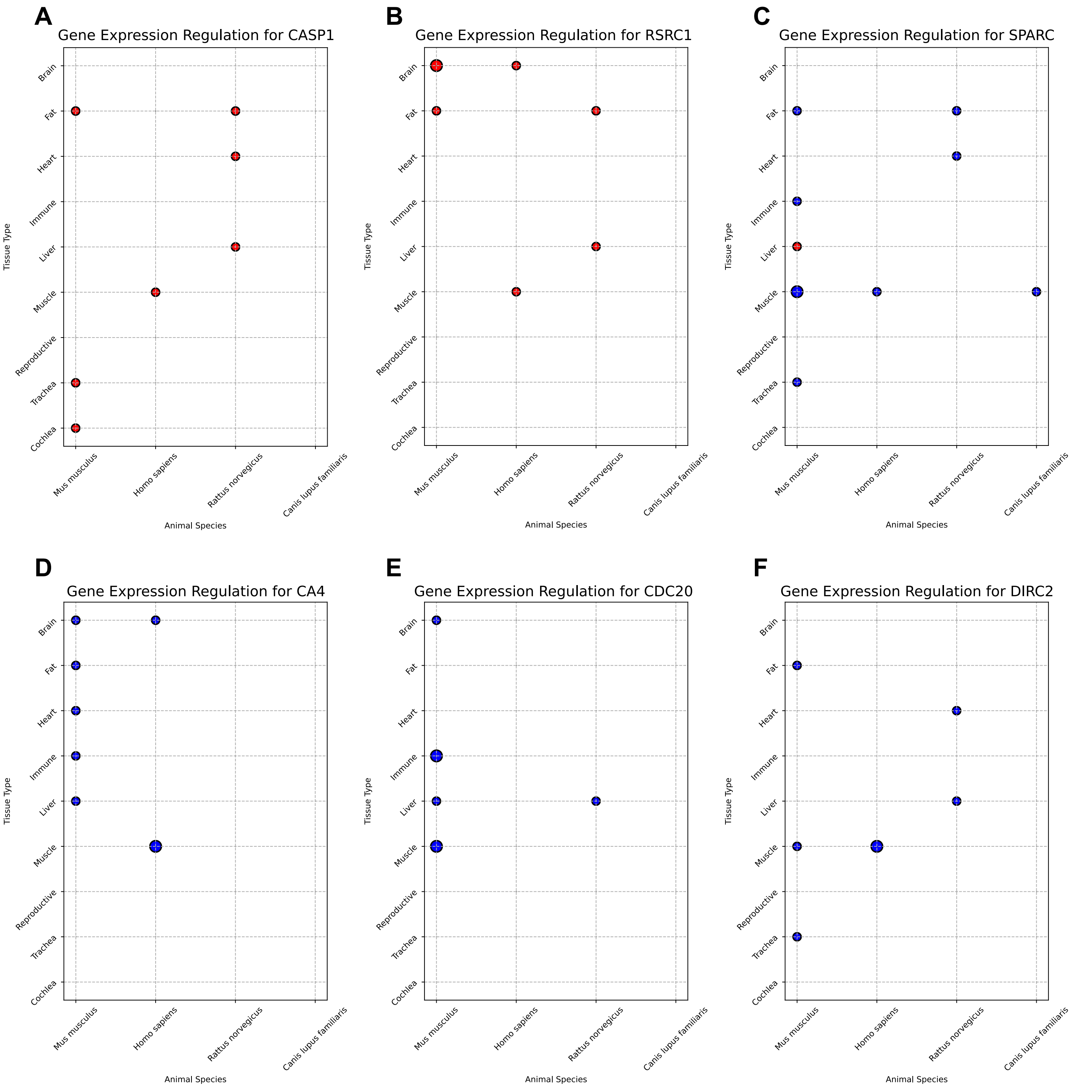


**Supplementary Figure 2.** Bubble plots showing the expression patterns of the six drivers of aging identified in this study: CASP1, RSRC1, SPARC, CA4, CDC20, and DIRC2. Bubbles represent significant differential expression between young and old samples: color indicates direction (red for age-upregulated, blue for age-downregulated), and size of the circle indicates the number of datasets exhibiting the displayed trend. Tissue of origin is plotted on each Y-axis (brain, fat, heart, immune, liver, muscle, reproductive, trachea, and cochlea), and species of origin is plotted on each x-axis (mouse, human, rat, dog).
